# Community-driven data analysis training for biology

**DOI:** 10.1101/225680

**Authors:** Bérénice Batut, Saskia Hiltemann, Andrea Bagnacani, Dannon Baker, Vivek Bhardwaj, Clemens Blank, Anthony Bretaudeau, Loraine Brillet-Guéguen, Martin Čech, John Chilton, Dave Clements, Olivia Doppelt-Azeroual, Anika Erxleben, Mallory Ann Freeberg, Simon Gladman, Youri Hoogstrate, Hans-Rudolf Hotz, Torsten Houwaart, Pratik Jagtap, Delphine Larivière, Gildas Le Corguillé, Thomas Manke, Fabien Mareuil, Fidel Ramírez, Devon Ryan, Florian Christoph Sigloch, Nicola Soranzo, Joachim Wolff, Pavankumar Videm, Markus Wolfien, Aisanjiang Wubuli, Dilmurat Yusuf, Galaxy Training Network, Rolf Backofen, James Taylor, Anton Nekrutenko, Björn Grüning

## Abstract

The primary problem with the explosion of biomedical datasets is not the data itself, not computational resources, and not the required storage space, but the general lack of trained and skilled researchers to manipulate and analyze these data. Eliminating this problem requires development of comprehensive educational resources. Here we present a community-driven framework that enables modern, interactive teaching of data analytics in life sciences and facilitates the development of training materials. The key feature of our system is that it is not a static but a continuously improved collection of tutorials. By coupling tutorials with a web-based analysis framework, biomedical researchers can learn by performing computation themselves through a web-browser without the need to install software or search for example datasets. Our ultimate goal is to expand the breadth of training materials to include fundamental statistical and data science topics and to precipitate a complete re-engineering of undergraduate and graduate curricula in life sciences.

## Introduction

Rapid development of DNA sequencing technologies has made it possible for biomedical disciplines to rival the physical sciences in data production capability. The combined output of today’s sequencing instruments has already surpassed the data generation speed of resources such as the Large Hadron Collider and is rivaling those in the field of astronomy. Yet biology is different from physics (and other quantitative disciplines) in one fundamental aspect—the lack of computational and data analysis training in standard biomedical curricula. Many biomedical scientists do not possess the skills to use or even access existing analysis resources. Such paucity of training also negatively impacts the ability of biological investigators to collaborate with their statistics and math counterparts, because of the inability to speak each other’s language. In addition, an estimated one-third of biomedical researchers do not have access to proper data analysis support [1]. The only way to address these deficiencies is with training. The need for such training cannot be overstated: while the majority (>95%) of researchers work or plan to work with large datasets, most (>65%) possess only minimal bioinformatics skills and are not comfortable with statistical analyses [1,2]. This overwhelming need drives the demand, which, at present, greatly exceeds supply [3]. In a recent survey [4] over 60% of biologists expressed a need for more training while only 5% called for more computing power. Thus one can assume that the true bottleneck of the current data deluge is not storage or processing power, but the knowledge and skills to utilize *existing* resources and tools.

Since 2006 our team has been pondering the question of how to enable computationally naive users to perform complex data analysis tasks. We attempted to solve this problem by creating a platform, Galaxy (http://galaxyproject.org [5]), that provides access to hundreds of tools used in a wide variety of analysis scenarios. It features a web-based user interface while automatically and transparently managing underlying computation details. It can be deployed on a personal computer, heterogeneous computer clusters, as well as computation systems provided by Amazon, Microsoft, Google and other clouds such as those running OpenStack. Over the years a community has formed around this project, providing it with an ever-growing, up-to-date set of analysis tools and expanding it beyond life sciences.

These features of Galaxy attracted many biomedical researchers, making it well suited for use as a teaching platform. Here we describe a community-driven effort to build, maintain, and promote a training infrastructure designed to provide computational data analysis training to biomedical researchers worldwide.

## Results and Discussion

Our goal is to develop an infrastructure that facilitates data analysis training in life sciences. At a minimum it needs to provide an interactive learning platform tuned for current datasets and research problems. It should also provide means for community-wide content creation and maintenance, and, finally, enable trainers and trainees to use the tutorials in a variety of situations such as those where a reliable Internet access is not an option.

### Interactive learning tailored to research problems

We produced a collection of hands-on tutorials that are designed to be interactive and are built around Galaxy. The hands-on nature of our training material requires that a trainee has two web browser windows open side-by-side: one pointed at the current tutorial and the other at a Galaxy instance. We build most tutorials around a “research story”: a scenario inspired by a previously published manuscript or an interesting dataset (with the caveat that some more technical materials do not lend themselves to this goal). To make training comprehensive, we aim to cover major branches of biomedical big data applications such as those listed in Table 1.

**Table 1.** Topics available in the Galaxy training material website (https://training.galaxyproject.org) with their target users and available tutorials. Admin = Galaxy administrators, Biol = Biomedical researchers, Dev = Tool and software developers, Inst = Instructors and Tutorial developers. The scripts for the extraction of such information are available in GitHub (https://github.com/bebatut/galaxy-training-material-stats). This table displays content current as of 28th Sep, 2017.

| Topic | Target | Tutorials |
| --- | --- | --- |
| Galaxy Server administration | Admin | Galaxy Database schema, Docker and Galaxy, Advanced customisation of a Galaxy instance |
| Assembly | Biol | Introduction to Genome Assembly, De Bruijn Graph Assembly, Unicycler Assembly |
| ChIP-Seq data analysis | Biol | Identification of the binding sites of the T-cell acute lymphocytic leukemia protein 1 (TAL1), Identification of the binding sites of the Estrogen receptor |
| Development in Galaxy | Dev | Contributing with GitHub, Tool development and integration into Galaxy, Tool Shed: sharing Galaxy tools, Galaxy Interactive Tours, Galaxy Interactive Environments, Visualizations: charts plugins, Galaxy Webhooks, Visualizations: generic plugins, BioBlend module, a Python library to use Galaxy API, Tool Dependencies and Conda, Tool Dependencies and Containers, Galaxy Code Architecture |
| Epigenetics | Biol | DNA Methylation |
| Introduction to Galaxy | Biol | Galaxy 101, From peaks to genes, Multisample Analysis, Options for using Galaxy, IGV Introduction, Getting data into Galaxy |
| Metagenomics | Biol | 16S Microbial Analysis with Mothur, Analyses of metagenomics data - The global picture |
| Proteomics | Biol | Protein FASTA Database Handling, Metaproteomics tutorial, Label-free versus Labelled - How to Choose Your Quantitation Method, Detection and quantitation of N-termini via N-TAILS, Peptide and Protein ID, Secretome Prediction, Peptide and Protein Quantification via Stable Isotope Labelling (SIL) |
| Sequence Analysis | Biol | Quality Control, Mapping, Genome Annotation, RAD-Seq Reference-based data analysis, RAD-Seq de-novo data analysis, RAD-Seq to construct genetic maps |
| Train the trainers | Inst | Creating a new tutorial - Writing content in markdown, Creating a new tutorial - Defining metadata, Creating a new tutorial - Setting up the infrastructure, Creating a new tutorial - Creating Interactive Galaxy Tours, Creating a new tutorial - Building a Docker flavor for a tutorial, Good practices to run a workshop |
| Transcriptomics | Biol | De novo transcriptome reconstruction with RNA-seq, Reference-based RNA-seq data analysis, Differential abundance testing of small RNAs |

As an example, suppose that a researcher is interested in learning about metagenomic data analyses. The category “Metagenomics” at https://training.galaxyproject.org presently contains a set of introductory slides, two hands-on tutorials (Fig. 1A), and HTML-based slides designed as a brief (10 - 20 min) introduction to the subject. In addition, every hands-on tutorial begins with background information (Fig. 1B) and explains how it influences data analysis. This background story is included to account for situations when tutorials are used for self-teaching in the absence of an instructor who would provide a formal introduction. After the introduction, the hands-on part of the tutorial begins and is laid out in a step-by-step fashion with explanations (boxes on Figure 1D) of what is being done inside Galaxy, which parameters are critical, and how modifying parameters affects downstream results. The first step in this progression is usually a description of the datasets and how to obtain them. We invested a large effort in creating appropriate datasets by downsampling original published data, which is necessary since real-world datasets are usually too big for tutorials. Our goal was to make datasets as small as possible while still producing an interpretable result. We use Zenodo (http://www.zenodo.org), an open data archiving and distribution platform, to store the tutorial datasets and to provide them with stable digital object identifiers (DOIs) that can be used to credit their authors and for citation purposes.

**Figure 1.**
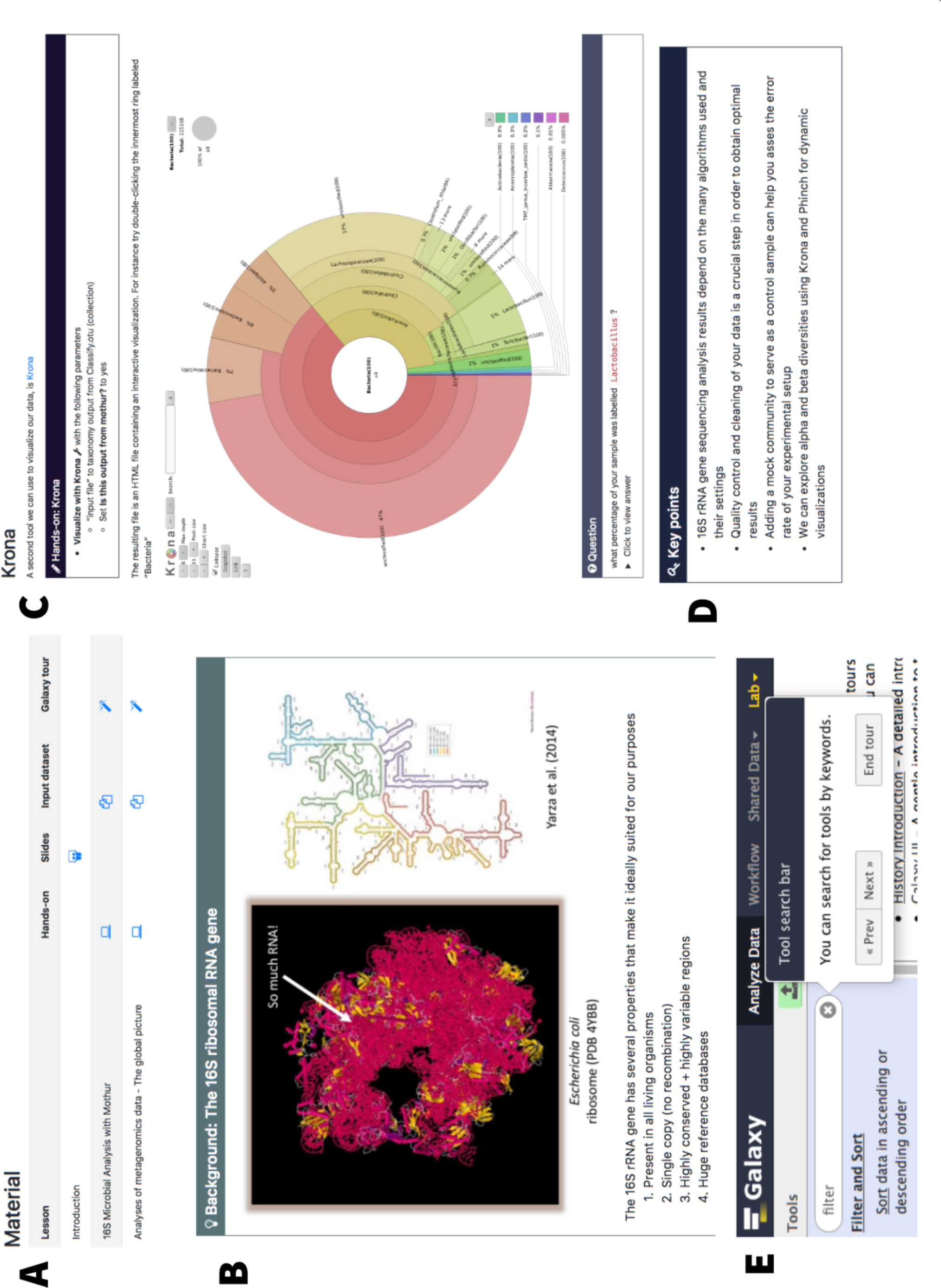

Tutorials start with a list of prerequisites (typically other tutorials within the site) to account for the variation in trainees’ backgrounds, a rough time estimate, questions addressed during the tutorial, learning objectives, and key points (e.g., Fig. 1D). These components help trainees and instructors to keep track of the training goals. For example, the learning objectives are single sentences describing what a trainee will be able to do as a result of the training [6]. Throughout the tutorials, question boxes (Fig. 1C) are added as an effective way to motivate the trainees [7,8] and guide self-training. The training material is distributed under a CC BY 4.0 (https://creativecommons.org/licenses/by/4.0/) license: its contents can be shared and adapted freely as long as appropriate credit is given. Efforts have been made also in the direction of ensuring website accessibility to disabled persons by regular evaluation with WAVE (http://wave.webaim.org), a web accessibility evaluation tool, and by automatic checking for alternative text for the images.

Keeping trainees engaged is critical, particularly for self-training. To this end, we aim to provide interactive tours for each tutorial: using instruction bubbles (Fig. 1E), each tutorial step can be “played” directly inside Galaxy, guiding learners to the needed tools while also allowing exploration of the framework’s functionalities. Tours can be created directly in the browser using our tour creator plugin (https://zenodo.org/record/830481).

### Infrastructure to facilitate community-led content development

To build a comprehensive collection of training materials covering the spectrum of topics in the life sciences, we must leverage community expertise, as no single group can possibly “know it all”. To achieve this goal, we built an infrastructure that makes tutorial creation a convenient, hassle-free process and enables transparent peer-review and curation to guarantee high-quality and current content. In implementing these requirements, we took inspiration from the Software and Data Carpentry [9] projects (SDC). In SDC, materials are openly reviewed and iteratively developed on GitHub (https://github.com/) to capture the breadth of community expertise. SDC delivers training via online tutorials with hands-on sections, which offer better training support than videos because trainees who are actively participating learn more [7]. This format is also adapted to face-to-face courses and self-training, as the content is openly accessible online. The content of these web pages is easy to edit, thus reducing the contribution barrier. The tutorials are developed in Markdown, a plain text markup language, which is automatically transformed into web-browser accessible pages. Using these strategies, we created a GitHub repository (https://github.com/galaxyproject/training-material) to collect, manage, and distribute training materials. The architecture of this infrastructure is shown in Fig. 2 (center), with the process for developing a tutorial illustrated at the bottom of the figure. To create a new tutorial, the main repository is *forked* (duplicated into a user-controlled space) within GitHub by an individual developing the tutorial. The developer then proceeds to write the content using Markdown as explained in our guide at https://training.galaxyproject.org/topics/training (itself consisting of several tutorials). The guide contains detailed information on technical and stylistic aspects of tutorial development. After settling on a final version of the tutorial (circles 1 through 10, the bottom of Fig. 2), a *pull request* is created against the original repository. When a new pull request is issued, this is an indication that a new tutorial is ready to be reviewed by the editorial team. The team then makes suggestions on the new contents, these suggestions are discussed, and the content is edited accordingly. A decision is then made whether to accept the pull request. At the same time the pull request is first created, the newly added content is automatically tested for HTML generation and all links and images are verified. When the pull request is accepted, the new tutorial becomes a part of the official training material portfolio, and the entire site is regenerated. This open strategy for content creation started paying off early as we already have over 60 individuals contributing and editing content within the GitHub repository.

**Figure 2.**
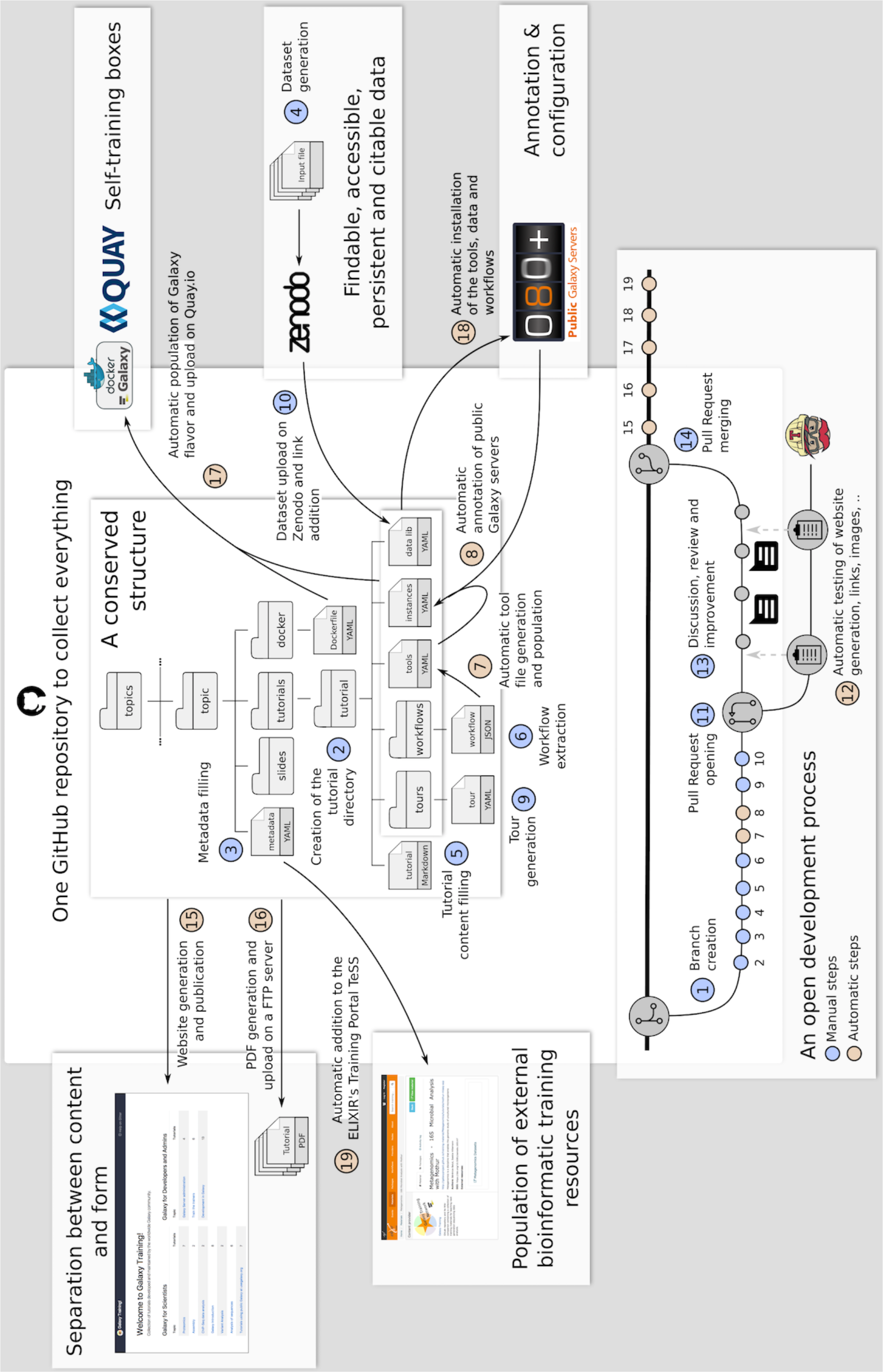

This infrastructure has been developed in accordance with the FAIR (Findable, Accessible, Interoperable, Reusable) principles [10]. Each tutorial, slide deck, and topic is complemented by numerous metadata described in a standard, accessible, interoperable format (YAML; http://yaml.org/). The metadata is used to automatically populate the TeSS training portal at the European life-sciences Infrastructure for biological Information (ELIXIR; https://tess.elixir-europe.org), ensuring global reach [11]. Each topic, tutorial, and slide deck has as metadata a reference to a topic in the EDAM ontology [12], a comprehensive catalog of well-established, familiar concepts that are prevalent within bioinformatics and computational biology. These references can be used to represent relationships among the materials and make them more findable and searchable.

Using the framework described above, we relaunched the Galaxy Training Network (GTN; https://galaxyproject.org/teach/gtn). This growing network currently consists of 33 scientific groups (https://galaxyproject.org/teach/trainers) invested in Galaxy-based training. The GTN regularly organizes training events worldwide (Fig. 3) and offers best practices for developing Galaxy-based training material, advice on compute platform choice to use for training, and a catalog of existing training resources for Galaxy (Table 1).

**Figure 3.**
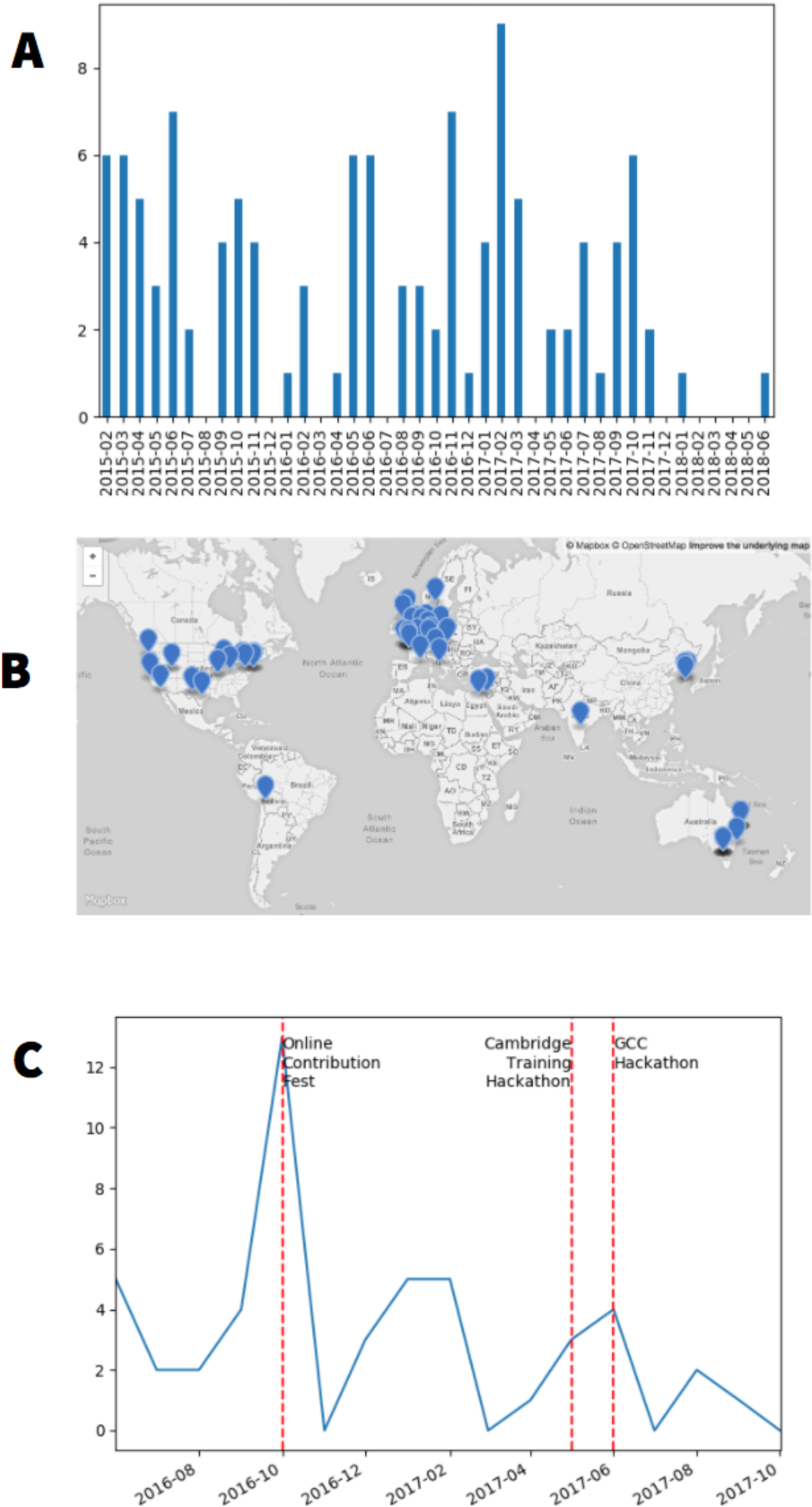

### Ensuring accessibility of tutorials

Most training materials hosted within the GTN resource are intended to be used side-by-side with the Galaxy framework. However, the main public Galaxy instances (e.g. https://usegalaxy.org or https://galaxy.uni-freiburg.de) are occasionally subject to unpredictable load, may be inaccessible due to network problems in remote parts of the world or may not have all the tools necessary for completing the tutorials. To account for these situations, we have developed a Docker-based framework for creating portable, on-demand Galaxy instances specifically targeted for a given tutorial. Docker (https://www.docker.com) is a container platform which provides lightweight virtualization by executing "images" (files that include everything needed to run a piece of software) isolated from the host computer environment. An individual creating a new tutorial lists all tools that are required to complete it in a dedicated configuration file (tools file, Fig. 2). For example, a metagenomics tutorial uses the mothur [13] set of tools as well as visualization applications such as Krona [14]. The corresponding Galaxy tools are listed in a configuration file that is a part of the metagenomics tutorial. This file is used to install these Galaxy tools and their dependencies into a base Galaxy Docker image (containing essential Galaxy functionality and a core set of tools) to create a dedicated “on-demand” Galaxy instance which can then be used on any trainer’s or trainee’s computer. The Docker image also contains input data, tours, workflows.

### A vision for the future

Life sciences are on a trajectory towards becoming an entirely data-driven scientific domain. A growing understanding that biomedical curricula must be modernized to reflect these changes is gaining attention [15]. Our project represents one of the first fully open, “grass-roots” attempts at unifying and standardizing heterogeneous training resources around the Galaxy platform. While it may not be appropriate to all, our multi-year experience with teaching workshops at various skill-levels can be summarized as the following set of recommendations, which we use as guiding principles. These recommendations may also be useful for the development of alternative frameworks as well as for curriculum planning:

1. **Require quantitative training.** No one expects biomedical researchers to rival their colleagues in departments of mathematics or statistics. However, background level statistical reasoning must be included in all training materials and general statistical courses must become a part of undergraduate and graduate education. This would have an enormous positive impact on the quality of biomedical research because researchers with basic understanding of quantitative concepts will not, for example, perform an RNAseq experiment without a sufficient number of replicates.
2. **Demystify computational methodologies.** Fundamental principles, limitations, and assumptions of molecular experimental techniques are typically well understood by biomedical researchers even when proprietary reagent kits are used. This is not the case with software tools, which are often treated as black boxes. We argue that fundamental principles of bioinformatic techniques (e.g., read mapping, read assembly) must be understood by experimentalists as this will also lead to an increase in overall quality of research output.
3. **Advocate the fundamental virtues of open and transparent research.** Open and transparent data analysis (e.g., through the use of open-source software) promotes replication and validation of results by independent investigators. It also speeds up research progress by facilitating reuse and repurposing of published analyses to different datasets or even to other disciplines. We advocate openness as a basic principle for computational analysis of biomedical data.

The infrastructure presented here has been developed to support training using Galaxy, a powerful tool for teaching bioinformatics concepts and analysis. But such a model is not only limited to Galaxy. It could be applied to bioinformatics training more generally (and to other disciplines as well) to support learners and instructors in this ever-changing landscape that is the life sciences.

## Acknowledgments

The authors are grateful to the Freiburg Galaxy and Core Galaxy teams, as without these resources this work would not be possible. Adoption of Galaxy Tours has been accelerated with the introduction of Galaxy Tour Builder (https://zenodo.org/record/830481) by William Durand (https://tailordev.fr). This project was supported by Collaborative Research Centre 992 Medical Epigenetics (DFG grant SFB 992/1 2012), German Federal Ministry of Education and Research (BMBF grant 031 A538A RBC (de.NBI)), NIH Grants U41 HG006620 and R01 AI134384-01, as well as NSF Grant 1661497.

